# Ethologically relevant behavioural assay for investigating reach and grasp kinematics during whole-body motor control in mice

**DOI:** 10.1101/2025.04.04.647225

**Authors:** Christopher J. Black, Liam E. Browne, Robert M. Brownstone, Stephanie C. Koch

## Abstract

Reach and grasp are critical components of skilled mammalian motor control and their detailed analysis in rodents has been key to deepening our understanding of prehension in the context of health and disease. However, most studies investigating these behaviours focus on isolating forelimb movements with little regard to the whole-body movements that are key for effective behaviour. To address this issue, we designed a novel behavioural approach to investigate reach and grasp during whole-body, vertical locomotion in mice. Using a customizable transparent climbing surface, we show that our behavioural approach can extract key kinematic features of climbing. Mouse climbing gait reflects aspects of quadrupedal locomotion, showing similar phase dependencies on increasing speed, including reduced stance (i.e. grasp) time and duty factor. Analysis of multi-limb coordination indicated that climbing revolves around anti-phasic forepaw movements with less consistency in interlimb coordination in the hindpaws. Fore- and hindpaws also differed in their reach trajectories and velocity profiles. The flexibility of this approach also allows for tailored climbing configurations, which we use to show that mice can adapt to and overcome vertical obstacles. By leveraging naturalistic climbing, our modular behavioural approach enables investigation of complex prehensile behaviours and facilitates new study into the neural circuits underlying whole-body skilled motor control.

## Introduction

It takes a symphony of motor behaviours to interact with the world. We fluidly combine walking and object manipulation when grabbing coffee, coordinate synchronous and asynchronous hand and foot movements to drive into work, and employ infinitely complex motor patterns during challenging tasks such as downhill skiing, playing the piano, and bouldering. Such skilled behaviours are composed of several fundamental sub movements, such as reach and grasp.^1,2^ Importantly, reaching and grasping are critical for both interacting with and manipulating objects, and are predicated on rapid decision making during grasp and fine tuning of motor control with continuous feedback from internal and external sensory information.^3^ Understanding the neural mechanisms underlying skilled reaching and grasping is thus a major area of focus in systems, computational, and translational neuroscience.

Advances in rodent models of skilled forelimb behaviours have allowed us to gain more insight into the neural mechanisms of reaching and grasping kinematics. Forelimb joystick and reach-to-grasp paradigms for mice have been used to investigate how the central nervous system regulates motor control and adaptation.^4–7^ Naturalistic reach and grasp behaviors, such as the pellet grasp, have also been used to examine deficits in skilled motor control in neurodegenerative conditions.^8–10^ However, current forelimb behaviors in rodents prohibit or minimize movement of the rest of the body. Given that reaching and grasping are fundamental components to many complex motor behaviors, these experimental approaches make it unclear how prehensile kinematics are embedded within whole-body movements.

One way in which mice naturalistically engage in reach and grasp during whole-body movement is during climbing. Mice are expert climbers; their ability to ascend vertical surfaces provides an additional spatial dimension in which they can forage for food, avoid predators, and explore their environment.^11^ Climbing is a unique behavior that combines quadrupedal locomotion,^12^ whole-body coordination and skilled multi-limb reaching and grasping,^13^ postural control,^14^ and many other features of complex motor control. In rodents, climbing behaviours have been used to evaluate motor deficiency during traumatic brain injury,^15^ pain,^16^ and neurodegeneration.^17^ However, obstruction of limb movements during climbing restricts quantitative analyses of climbing to duration, distance, and speed. To overcome this issue, we have developed a novel climbing assay that leverages a unique transparent climbing wall and markerless pose estimation to extract paw kinematics during naturalistic climbing. We show that mice display an innate drive and ability to climb on this novel substrate, without the need for training, and can even overcome vertical obstacles. Our results show that climbing kinematics reflect aspects of both horizonal locomotion and whole-body coordination centered around forelimb reach and grasp, allowing for a novel and ethologically relevant behavioural paradigm for the study of sensorimotor neural circuits, navigation and learning.

## Results

### Modular Climbing Rig for Automated Ventral Imaging of Paw Kinematics

To examine paw kinematics during climbing, we designed a modular behavioral rig to perform high-speed imaging (200fps) of the ventral surface of the mouse through a transparent climbing wall (Figure 1A and 1B). SLEAP^18^ was used to perform markerless pose estimation to track the nose, base of the tail, and paws during climbing (Figure 1B). A force plate in the base of the enclosure and a motion sensor at the top of the climbing wall to automate high-speed video recordings (Figure 1C) by employing the Bonsai framework.^19^

**Figure 1.**
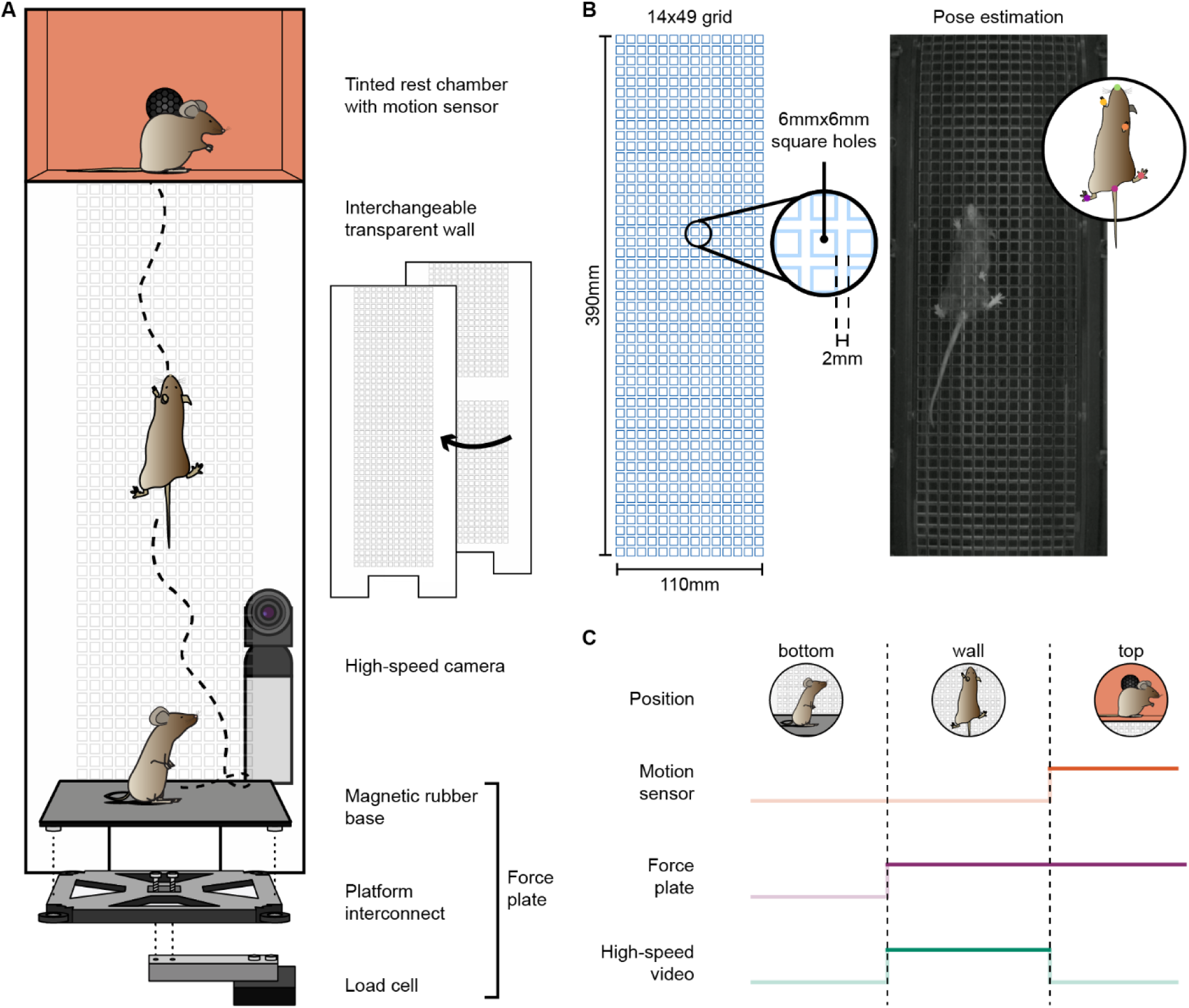
Schematic of climbing behavior. (A) Climbing apparatus consists of a custom force plate to track animals moving from the base to the wall, interchangeable transparent climbing wall, and resting chamber with motion sensor to detect successful climbs. A high-speed camera behind the climbing wall captures the ventral surface of the mouse during climbing. (B) Dimensions of the climbing grid layout (left) with an example frame extracted from a video (right) of a climbing mouse for use in markerless pose estimation. (C) Sensor based video acquisition control scheme.

Prior to behavioral testing, we allowed mice to habituate to the behavioral rig for a minimum of three days. Mice are known to perform spontaneous climbing in their home cage^20^; similarly, we found that mice exhibited spontaneous wall climbing during habituation (Figure S1). During habituation, 80% of mice spontaneously climbed (2.4± 2.0 climbs/session). To increase the number of trials performed during behavioural testing, mice were gently coaxed towards the wall by lightly tapping their tail with a soft brush.

### Repetition does not alter climbing performance

Skilled motor performance is known to improve with repetition.^21^ As climbing is an innate and highly skilled behavior, we sought to determine whether performance metrics changed over repeated climbing sessions. To quantify motor performance, we examined climbing speed and paw grasping accuracy. Overall speed for each trial was defined by the time it took to traverse a 20cm area of the wall (Figure 2A). This area was selected to account for differences in vertical starting position between trials. We compared the average traversal speed across testing sessions (Figure 2B) and found no significant difference across sessions (p=0.98). Accuracy was examined by quantifying the number of paw slips (Figure 2C).^22^ Similar to average speed, the average number of paw slips did not change significantly across sessions (p=0.99). These results show that motor performance remains consistent across sessions.

**Figure 2.**
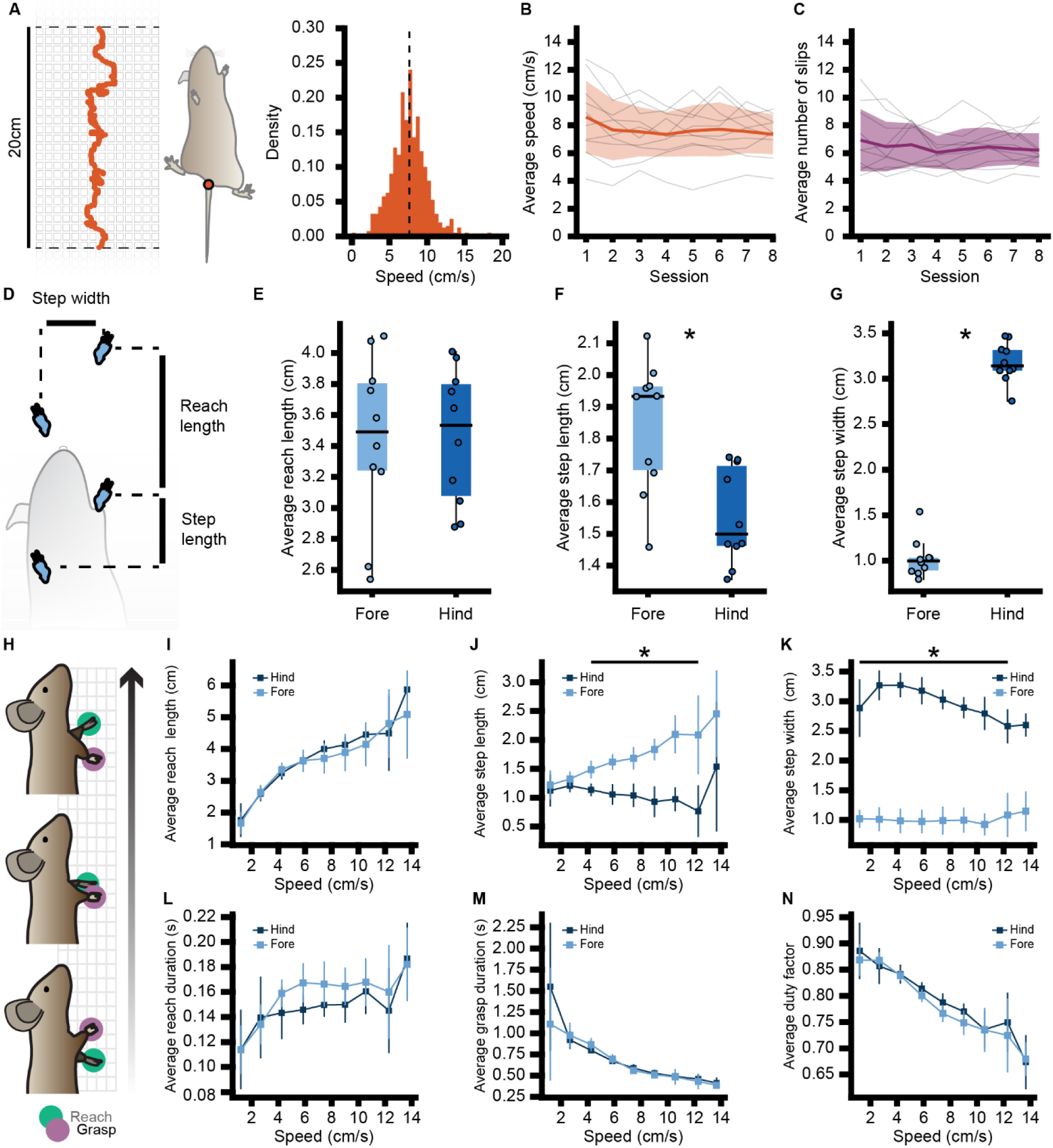
Climbing gait features alter with speed. (A) Trace of tail base (orange) for one climb within 20cm region for calculating climbing speed (left). Histogram of speed (right) across all sessions was 7.65±2.41cm/s (800 climbs, 10 mice). (B) Average speed across 8 testing sessions for all mice (n=10, p=0.98, Kruskal-Wallis). (C) Average number of slips for each paw across 8 testing sessions for all mice (n=10, p=0.99, Kruskal-Wallis). (D) Gait parameters applied to vertical climbing to examine reach length, step width, and step length. Boxplots with individual data points showing forepaw (light blue) and hindpaw (dark blue) values for (E) average reach length, (F) average step length, and (G) average step width (n = 10, *p < 0.05, Wilcoxon). (H) Illustration of kinematic variables for climbing gait by accounting for vertical speed (I-K) Comparison of same gait features as in (E-G) but averages are placed in 5cm/s bins based on corresponding body speed. (L) average reach duration, (M) average grasp duration, and (N) average duty factor for fore (left) and hind (right) paws (n=10 mice).

### Speed affects vertical gait

We next characterized vertical analogs of horizontal gait features (Figure 2D) such as fore- and hindpaw reach length (vertical distance between consecutive grasps), step length (vertical distance between opposing paws), and step width (horizontal distance between opposing paws). We found that mice displayed similar reach lengths between forepaws (3.4±0.5cm) and hindpaws (3.5±0.4cm). However, step length was significantly greater (p < 0.05) for forepaws (1.8±0.2cm) than hindpaws (1.6±0.1cm), while step width was significantly smaller (p < 0.05) for forepaws (1.02±0.20cm) than hindpaws (3.18±0.21cm).

As speed has been shown to influence stepping and the duration of swing and stance phases of the gait cycle during horizontal locomotion,^23–25^ we hypothesized that the same would be true of vertical locomotion. To test this, we first calculated the calculated correlation coefficients for individual gait features with respect to climbing speed (Figure 2D-2H and Figure S2). There was a moderate (>0.30) positive correlation between speed and reach length (forepaw: 0.32±0.02, hindpaw:0.42±0.02), and a moderate (<-0.30) negative correlation between speed and duty factor (forepaw: −0.43±0.03, hindpaw:- 0.39±0.02) and speed and grasp phase (forepaw: −0.49±0.03, hindpaw:-0.43±0.04) for fore- and hindpaws. Intuitively, these results show that as climbing speed increases, mice spend less time grasping. These results are also analogous to changes in gait during horizontal locomotion, where higher velocity corresponds to decreased time in the stance phase.^23^ Although some features displayed weak to no correlation with speed (Figure S2), we wanted to determine if speed affected differences between fore- and hindpaw gait features. Therefore, we calculated the average gait features for reach length, step length, step width, grasp and reach duration, and duty factor for the fore- and hindpaws of each animal in 5cm/s bins (Figure 2I-2N). Fore- and hindpaws displayed no differences between average reach length, reach duration, grasp duration, or duty factor with respect to speed. Forepaws had significantly greater (p < 0.05) average step length than hindpaws at speeds between 4 and 12cm/s (Figure 2J), which reflects the difference in correlation between step length and speed (forepaw: 0.16±0.02, hindpaw: −0.05±0.01). Average step width was significantly greater (p < 0.05) for hindpaws than for forepaws at speeds below 12cm/s (Figure 2K). However, correlations between average step width and speed (forepaw: −0.03±0.01, hindpaw: −0.05±0.01) suggests fore- and hindpaw width is stable across different climbing speeds.

### Climbing revolves around anti-phasic forepaw coordination

As climbing requires inter and intrasegmental coordination, we next sought to examine how fore- and hindpaws coordinate during climbing. We will use previously defined terms to reference left and right forepaw or hindpaw pairs as homologous, forepaw and hindpaw pairs on the same side as homolateral, and forepaw and hindpaw pairs on opposite sides as diagonal (Figure 3A and 3B).^26^ To quantify coordination, we calculated the phase offset between paw pairings during reaching (Figure 3B).

**Figure 3.**
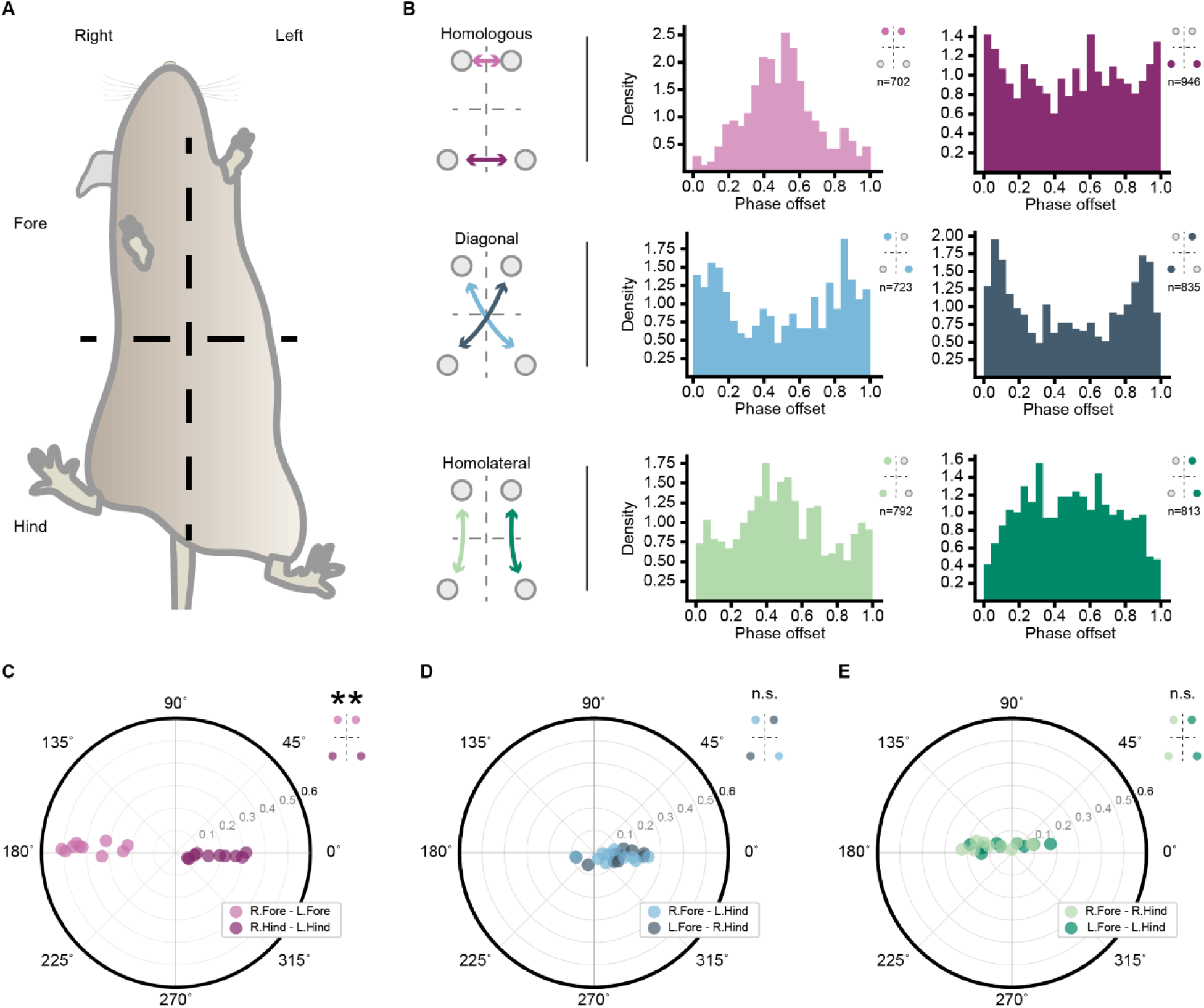
Climbing elicits modular coordination profile. (A) Mouse body axis for climbing illustrates limb references for ventral imaging. (B) Example histograms from one mouse of phase offset values between homologous (top left, purple) fore limb pairs (top middle) and hind limb pairs (top right); diagonal (middle left, blue) fore limb pairs (center) and hind limb pairs (middle right); and homolateral (bottom left, green) left fore and hind limb pairs (bottom middle) and right fore and hind limb pairs (bottom right). (C) Polar plots for homologous (left, purple), diagonal (middle, blue), and homolateral (right, green) paw pairs (light and dark shades specify limb groupings) show average phase preference in degrees and mean resultant vector for each animal (n=10, **p < 0.001, Watson-Williams).

Homolateral pairs exhibited an anti-phase relationship on left (124.56±61.90°; Rayleigh r = 0.41, p = 0.18) and right (128.99±54.73°; Rayleigh r = 0.54, p = 0.048) sides. Diagonal pairs displayed a significant in-phase relationship when comparing right forepaws and left hindpaws (342.53±39.48°, Rayleigh r = 0.76, p < 0.05) and left forepaws and right hindpaws (342.97±46.36°, Rayleigh r = 0.67, p < 0.05). Forepaw homologous pairs exhibited a strong anti-phase preference, akin to locomotion (176.40±3.50°, Rayleigh r = 0.99, p < 0.001), whereas hindpaws displayed an opposite in-phase preference akin to hopping (352.19±8.78°, Rayleigh r = 0.99, p < 0.001).

While mice displayed similar phase preferences within homolateral and diagonal pairings, homologous pairings were significantly different (p < 0.05) with forepaws favoring anti-phase coordination and hindpaws favoring weak in-phase coordination (Figure 3C). These results indicate that climbing revolves around alternating forepaw reaching and grasping with coordination between other paw pairings being less robust.

### Forepaw reach trajectories bias deceleration

Given that modulation of reach velocity is linked to successful completion of skilled point-to-point movements,^27^ we next sought to evaluate difference in the velocity of reach trajectories between fore- and hindpaws (Figure 4A). Reaches displayed a stereotypical bell-shaped velocity profile (Figure 4B) consisting of an acceleration and deceleration phase shown previously during rodent reach and grasp behaviors.^28^ Both forepaws and hindpaws spent more time in the deceleration phase than the acceleration phase (Figure 4B). While we found no significant difference in the average duration of the acceleration phase across all paws, both forepaws were found to have extended deceleration phases compared to hindpaws (p < 0.05) (Figure 4C and 4D). Increased deceleration phases in reach and grasp tasks may reflect increased accuracy demands for the given movement,^29^ suggesting that grasping accuracy during climbing is more important for forepaws than for hindpaws.

**Figure 4.**
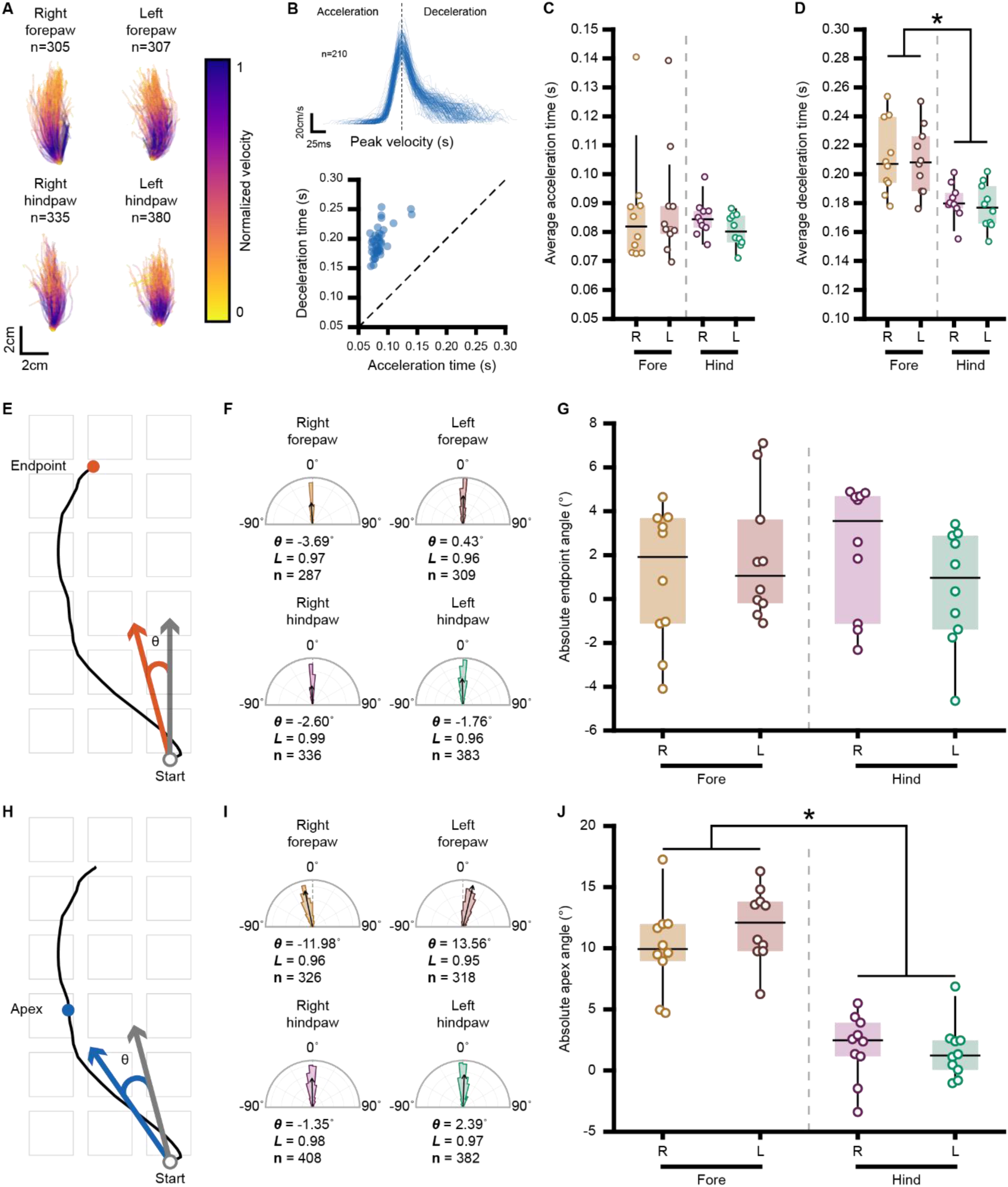
Fore limb reaching and grasping has higher skill requirement. (A) Example fore- and hindpaw reach trajectories from 80 trials from an individual mouse color coded to normalized speed. (B) Example forepaw reach and grasp velocity profiles (top) indicating acceleration and deceleration phases relevant for skilled point-to-point movements. Scatter plot of average time spent in acceleration phase versus deceleration phase (bottom). Each data point represents a single paw from one animal (n=40 paws from 10 mice), diagonal line through the origin indicates the bias of reach and grasp trajectories toward the deceleration phase. (C) Comparison of average time spent in acceleration phase and (D) deceleration phase across all paws for all mice (n=10, *p < 0.05, Kruskal-Wallis with Bonferroni correction). (E) Illustration of end point angle with (F) example polar distributions from one mouse. (G) Boxplots of average endpoint angles between all paws with individual data points (n=10 mice, p=0.48, Kruskal-Wallis). (H) Illustration of apex angle with (I) example polar distributions from one mouse. (J) Boxplots of average apex angles with individual data points (n=10, *p < 0.05, Wilcoxon with Bonferroni correction).

Given the freely moving aspect of climbing, we sought to determine whether the individual paw trajectories displayed any similarities with respect to reach directionality. To examine this, we first calculated tangent angles to the endpoint and apex of each trajectory with respect to the midline (Figure 4E-J). Doing so provided us with an estimate of the heading direction and curvature of the paw movement and the final target direction. For every animal we generated a distribution of endpoint angles from every trajectory for each paw (Figure 4F) to determine the preferred angle (Figure 4G). We found no significant difference between average endpoint angles or trajectory arcs for forepaws (endpoint angle: right: 0.99±3.13°; left: 1.91±2.95°; apex angle: right: 0.09±3.62°; left 11.87±3.02°, or hindpaws (endpoint angle: right: 2.32±2.91°; left: 0.53±2.63°; apex angle: right: 1.93±3.13 left: 1.55±2.28°). However, average apex angles were significantly greater (p < 0.05) for forepaw than hindpaw trajectories (Figure 4J). These results indicate that forepaw reaches produce more curvature in their trajectories than hindpaw reaches. As curvature is theorised to play a role in optimizing reach trajectories,^30^ this suggests that optimization of forepaw reaching is more necessary than hindpaw reaching during climbing.

### Mice employ a distinct grasping strategy for overcoming vertical obstacles

To highlight the modularity of the behavioural system and to examine the capability of mice to handle vertical obstacles, we designed an analog to the gap-cross task, in which mice need to use sensory information to locomote between separated platforms.^31,32^ This version of the climbing wall was based on the standard grid layout but featured a vertical 3cm (3.4 cm total) section of etched acrylic (Figure 5A) spanning the width of the wall that formed a tactile ‘gap’. We first examined whether gap crossing improved upon repeated testing (Figure 5A). We found that all mice were able to successfully cross the gap during the first session (n=10), and while the number of attempts decreased by the fourth day, there was no significant improvement in success rate, or in average time-to-cross over the 4 sessions (Figure 5B and 5C).

**Figure 5.**
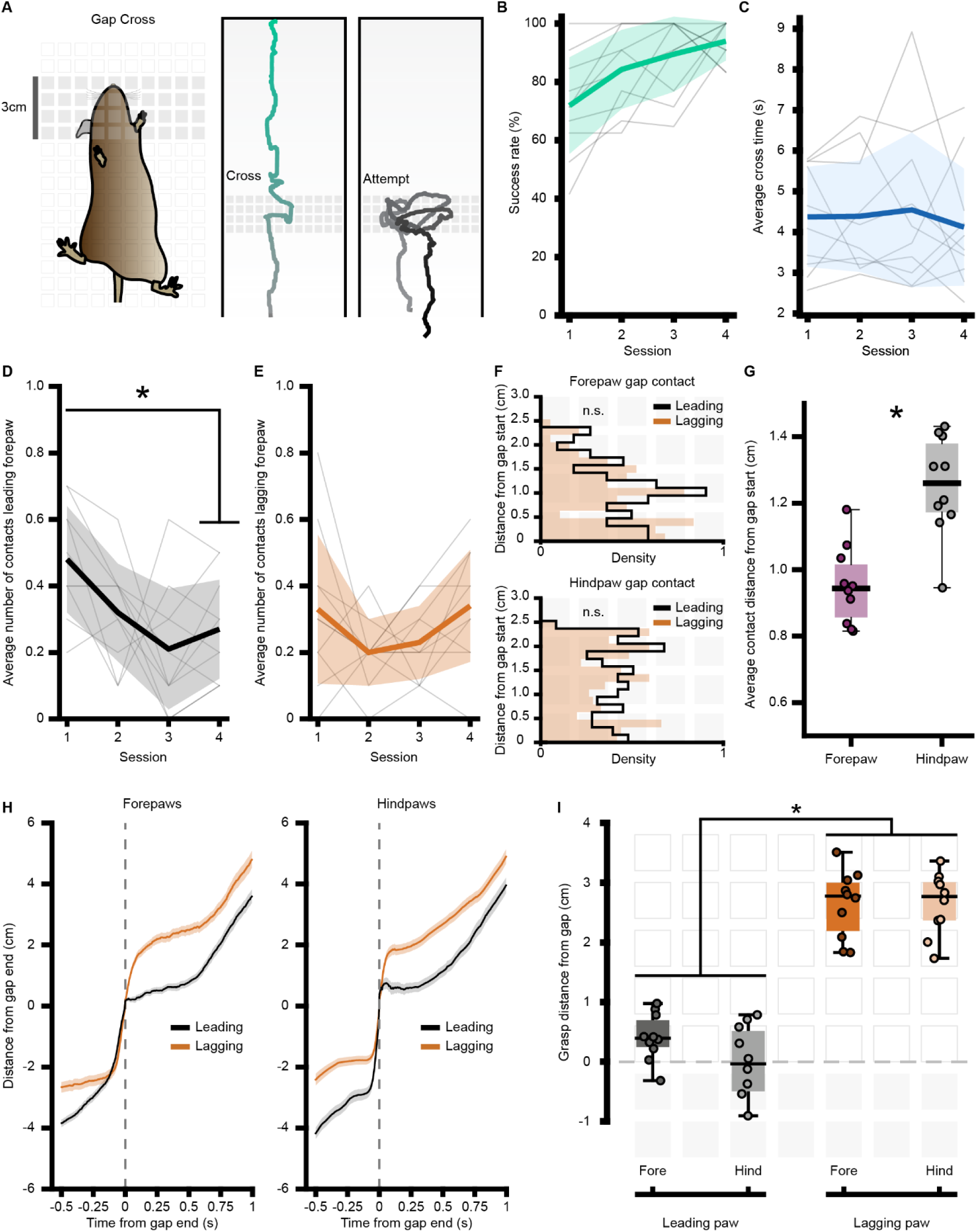
Vertical gap cross. (A) Illustration of the modified gap cross wall (left) with example traces from the nose of successful (center) and attempted (right) crosses. (B) Gap cross success rate grand average (green, mean ± standard deviation, n=10 mice) and individual averages (grey) across 4 sessions. (C) Gap cross time-to-cross grand average (blue, mean ± standard deviation, n=10 mice) and individual averages (grey) across 4 sessions. (D) Average number of gap contacts for the leading forepaw of all mice (orange, mean ± standard deviation) with individual averages (grey), (E) same as D) but for the lagging (black) forepaw. (n=10, *p < 0.05, paired t-test with Bonferroni correction). (F) Histograms showing contact location with respect to gap starting point for leading (black) and lagging (orange) forepaws (top), and hindpaws (bottom). (n.s., non-significant, two-sample Kolmogorov–Smirnov). (G) Boxplot of average gap contact position between grouped forepaw data (purple) and grouped hindpaw data (grey). (n=10, *p < 0.05, Wilcoxon). (H) Average vertical distance (mean ± 95% CI) from gap plotted with respect to time of gap crossing for leading (black) and lagging (orange) fore (left) and hind (right) paws. (I) Boxplot of average post-gap grasp positions for leading fore (dark grey) and hind (light grey) paws and lagging fore (dark orange) and hind (light orange) paws (n=10, *p < 0.05, Wilcoxon).

On some trials, mice would use their paws to contact the gap. We hypothesized that forepaw contact events could indicate a measure of learning, by which mice attempt to measure the start and end of the gap over time. To examine this, we first quantified the number of times the forepaws contacted the gap and then grouped the contacts by whether they occurred from the leading paw (i.e. the first paw to cross the gap) or the lagging paw (i.e. the second paw to cross the gap). We found that the average number of contact events of the leading forepaw significantly decreased (p < 0.05) from the first (0.48±0.16 contacts/trial) to last (0.27±0.15 contacts/trial) session (Figure 5D), a pattern not seen in the lagging forepaw (Figure 5E). These results suggest that the mice can optimize the leading forepaw movement across the gap over sessions, while the lagging forepaw movement remains unchanged. The hindpaws also contacted the gap during crossing, and we hypothesized that these contact points would differ from the forepaws based on the potentially reduced sensory information (i.e. visual and vibrissae input). To see whether there were any differences in these contacts, we looked at the positions at which the contacts were made. We first found that the position of gap contact did not differ significantly between leading and lagging pairs for either fore- or hindpaws (Figure 5F). We therefore grouped both the forepaw and hindpaw data and found the average contact position was significantly lower (p < 0.05) for the forepaws (0.95±0.11cm from the start of the gap) than the hindpaws (1.25±0.14cm) (Figure 5G).

The gap size forced mice to reach a minimum of ∼3.4cm to perform a successful cross, which was near the average reach length for fore- and hindpaws on the standard grid (Figure 2E and 2I). We therefore wanted to determine if this forced reach length altered the stepping, or grasp, length. To examine this, we looked at the vertical positioning of the paws with respect to the gap cross time (Figure 5H and 5I). We found that on average, leading paws grasped almost immediately after the gap (average forepaw position: 0.41±0.37cm; average hindpaw position: −0.18±0.89cm), while lagging fore- and hindpaws similarly grasped at significantly higher (p < 0.05) positions (average forepaw position: 2.63±0.53cm; average hindpaw position: 2.65±0.49cm). In comparison to step length on the standard grid (Figure 2F and 2J), there was no significant difference in step length between the fore- and hindpaws following the gap. These results suggest that the leading paw may play a support role for the lagging paw to increase the mouse’s distance from the gap.

## Discussion

Our novel behavioral platform extends the utility of climbing assays by introducing a way to extract limb kinematics through ventral imaging and markerless pose estimation. We quantitatively characterized reaching and grasping in the context of vertical locomotion and show that fore limbs display robust coordination that reflects a need for enhanced accuracy with respect to the hind limbs. By designing the system around a low cost, laser cut acrylic wall, we further extend the capabilities of our climbing assay by allowing users to rapidly prototype, test, and share different climbing substrates. In this respect, we show that mice can climb both a simple grid layout and cross a vertical ‘gap’. Our aim with this system is that it will help expand on prior assays that implement climbing to examine neural dynamics^33^ and evaluate disease states^16,20^ by providing insight into the neural mechanisms underlying complex whole-body movements.

We expected climbing performance (as measured by speed and paw slipping) to increase across testing days, however we instead found that performance was relatively stable (Figure 2B and 2C). One possible reason for this stability is that mice may already have reached their skill ceiling for climbing prior to behavioural testing. In the first postnatal month, climbing oriented harvest mice develop all the necessary elements for climbing,^13^ likewise, laboratory mice display climbing behaviour within this same period.^34^ This suggests that mice may already be well ’trained’ on climbing prior to behavioral testing, which may also explain why the more challenged gap cross task yielded a high success rate across all testing days. Therefore, behavioural testing during earlier developmental periods may allow the tracking of enhanced performance metrics during climbing. Another reason for stagnant performance is that the lack of external motivation for climbing might not drive any necessary improvement in behaviour. This was evident on some trials where mice would pause mid-climb before continuing to the top.

Planning and successful execution of reach and grasp is predicated on integration of sensory information primarily in the visual, proprioceptive, and tactile domains.^35,36^ Sensory feedback for reach and grasp is reflected in the bell-shaped velocity profiles,^37^ which was shown during climbing (Figure 4B). Furthermore, forepaw contacts during vertical gap crossing (Figure 5D-5G) reflect the need for tactile information to update reach and grasp behaviours. Importantly, other forms of sensory information have been shown to influence reaching and grasping in humans.^38^ Likewise in rodents, the vibrissae system is critical in sensorimotor integration and has been shown to aid in orientation and retrieval during reach-to-grasp^39^ and primes execution of whole-body gap cross movements.^32^ Although we did not explicitly track whiskers during climbing in this study, the elevated vertical positioning of the nose relative to the forepaws suggests that the whiskers could provide critical information for preparation and planning of reach and grasp.

Posture is also an important factor in executing reach and grasp strategies^40^ and affects exploration area.^5^ During climbing, mice exhibited wide hind limb stances and narrow fore limb positioning for most vertical speeds (Figure 2K). Importantly, a wider base of support can aid in postural adjustments needed to optimize reaching and grasping,^41^ suggesting that a potential optimal strategy for climbing in mice is to stabilize their trunk and lower extremities to enhance fore limb reaching and grasping. This may help explain the relatively strong preference of homologous forelimb alternation and weak hind limb synchronicity (Figure 3C), as the movement of the hind limbs may be restrained by producing support for forelimb coordination.

High-speed ventral imaging is key to extracting temporally resolved whole-body climbing kinematics. Although this approach provides an advance for analyzing climbing behaviors in a quantitative manner, one drawback is the difficulty of resolving certain body parts (i.e., elbow, knee, hip) in the ventral plane making it impractical to analyze complex joint dynamics. However, prior locomotor studies identify a simple solution for this issue by utilizing angled mirrors to capture both ventral and sagittal body planes during continuous behavior.^42^ Although outside of the scope for this current implementation, future modifications of our climbing system could enable measurements of joint angles and help produce 3D reconstructed limb trajectories during climbing.

We were able to show that mice did not require training to climb, as mice climbed either through self-initiation or gentle coaxing. While coaxing mice was an effective strategy for examining motor control prior studies have incentivized vertical climbing through food^33^ and water^43^ rewards. This means future studies could couple reward driven climbing with novel wall designs to examine reach and grasp kinematics during complex decision-making processes.

## Methods

### Animals

All experiments received review committee approval and were carried out at University College London on project license number 8884544. Procedures conformed to UK Home Office regulations and were performed in accordance with the UK Animals (Scientific Procedures) Act 1986. All mice used in this study were group housed, C57/BL6J background (sourced from Charles River), and received ad libitum food and water.

### Climbing assay

Mice were habituated for 15-30 minutes in a small, padded chamber for 3-5 days with access to the climbing wall to gauge their propensity for spontaneous climbing. Mice were placed in the bottom of the climbing enclosure and were allowed to explore. When mice climbed to the top chamber, they were placed back to the bottom of the enclosure within 2 minutes. During testing, mice either climbed spontaneously or, if spontaneous climbing did not occur within 1 minute, were gently coaxed with light taps from a paint brush to initiate climbing. This helped ensure repeated trials across sessions and reduce overall time mice spent performing behaviour. Mice completed single sessions of less than 30 minutes consisting of no less than 10 climbs per session. Mice underwent 4-5 sessions per week and completed a minimum of 8 sessions for the standard wall.

Following the completion of behavioural testing on the standard wall, mice completed four 30-minute sessions on the gap cross wall. For each session mice were manually coaxed to initiate wall climbing until 10 successful gap crosses were made, with no more than 25 attempts being attempted during any given session. On trials where mice did not cross the gap but their nose and or forepaws did, were considered attempted crossings.

### Behavioural rig

The main compartment of the climbing rig was a custom laser cut enclosure supported by 25mm rails on an aluminum breadboard (Thorlabs). A custom matte-black acrylic door and detachable magnetic wall (Perspex, Cutlaser Cut LTD) were used to easily place animals in the enclosure and clean the behavioral rig following use. 3mm transparent acrylic side walls (Perspex, Cutlaser cut LTD) were used to observe the animal during experimental sessions.

The base chamber of the climbing rig consisted of a custom 3D printed (Protolabs Network, 3D Hubs) compartment housing a removable magnetic base plate attached to a 1kg load cell (Adafruit, 4540). Output from the load cell was amplified and digitized using an analog-to-digital converter (HX711, SparkFun). Voltage changes generated by the loadcell were sent to an Ardunio microcontroller to be thresholded in order to identify when the mouse moved off of the platform.

The climbing wall consisted of a single 2mm thick acrylic sheet (Perspex) with a custom grid that was laser cut to a final size of 400×225mm. All laser cutting was performed by CutLaser Cut LTD. The grids were formed by laser cutting 6mm square holes evenly spaced by 2mm. The standard grid was a 14×49 arrangement, while the gap variant was created by etching, as opposed to cutting, a 4×49 section of the grid to produce a tactile surface that the mice could not grip. Illustrator files used for laser cutting are available at https://github.com/cjblack/TheWall.

A high-speed machine vision camera (1.6MP Blackfly S, FLIR) with a wide-angle lens (4mm UC Series Lens, Edmund Optics) was placed ∼37.5cm behind the wall and rotated 90 degrees for ventral imaging. Red (625nm) LEDs were positioned behind the camera angled ∼45 degrees to evenly illuminate the climbing wall. This provided optimal lighting for pose estimation while also reducing unnecessary strain on the animal’s vision due to their decreased sensitivity past the 600nm range.

The top chamber consisted of a custom transparent red acrylic container (Perspex, laser cut in house) designed to minimize light entry. At the back of the chamber was a motion sensor that sent a digital trigger to an Arduino whenever the animal entered the chamber.

Digital triggers generated by the Arduino tracking the load cell state and digital triggers directly from the motion sensor were sent to a second Arduino microcontroller that communicated with the Bonsai-Rx software to automate starting and ending high-speed video acquisition.

### Markerless pose estimation

High-speed video data was collected at 200fps and cropped to 600×1440 pixels. Prior to analysis, video data was corrected for rotational and wide-angle lens distortions. SLEAP was used to perform labeling, training, and inference for markerless pose estimation. 300 frames taken from 15 videos (20 frames/video) were labeled for the right and left fore- and hindpaws, base of the tail, and the nose. Labeled frames were then used to train a model with a UNet backbone using hyper parameters listed in Table S1. For the final model the mean error distance was 3.1 pixels (0.7mm), the mean Average Precision (mAP) was 0.88, and the mean Average Recall (mAR) was 0.90. Training and inference were performed on a high-performance computing cluster with either an NVIDIA A100, Tesla V100, or Tesla P100 GPU.

### Kinematic Analysis

#### Gait

Velocity for vertical gait analysis was calculated as the total distance traversed by the base of the tail during a given period, Δt. The base of the tail was selected as it most closely reflected movement of the torso and was not affected by individual movement of fore or hind limbs. Reaching and grasping was defined on every trial for each paw by first calculating the paw’s X and Y velocity. The velocities were then thresholded to find reach initiation events calculated as the time prior to when velocities transitioned above threshold, and grasp events calculated as the time after velocities transitioned below threshold. These time stamps were used for subsequent analyses for both gait and reach and grasp. Paw slipping was determined by similarly thresholding the paw velocity to identify periods where the velocity reached below −10 cm/s. This value was selected from identifying slipping events in videos to corresponding kinematic data.

Step length was calculated as the vertical (y) distance between homologous paw pairs at the time of a grasping event, stride was calculated as the vertical distance between two consecutive grasps for one paw, and step width (base of support) was calculated as the horizontal (x) distance between homologous paw pairs. Reach phases were calculated as the time between reach initiation and subsequent grasp, while grasp phases were conversely calculated as the time between grasp and subsequent reach initiation. Duty factor was calculated as the ratio of time spent in grasp over the entire time of the climbing ‘stride’, which was defined as the time between subsequent grasping events.

#### Directionality

To determine the directionality of trajectories, we used the inverse tangent function to calculate the angle from the y-axis given the initial coordinates (x_0_, y_0_) of the trajectory to the apex (x_1_, y_1_) of the trajectory.

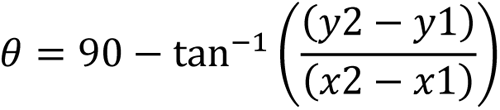

#### Coordination

To examine coordination, we utilized standard definitions for gait assessment. The climbing gait was determined by the time window in which each limb completed one ‘stride’. Strides for individual paws were determined by the period of successive wall contacts by the paw, which was determined by observing the rate of change in the X and Y coordinates. The phase offset was then calculated between inter- and intralimb paw pairings. Phase offset between a reference paw (P1) and a contralateral or ipsilateral paw (P1) is defined by adopting previous methodologies for calculating phase offset ^44^:

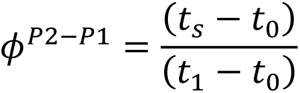

Where [t_0_, t_1_] gives the period between consecutive reach initiations of a reference paw P2, and t_s_ is the initiation reach time of another paw P1 during the reference period.

### Statistical testing

Normality was determined by the Shapiro-Wilks test for small sample sizes (n<50), with Kolmogorov-Smirnov test used for larger sample sizes, and the two-sampled Kolmogorov-Smirnov test used for comparing separate non-normal distributions of large sample sizes. Parametric comparisons were performed with a one-way ANOVA, with post-hoc paired t-test and Bonferroni corrections. Non-parametric comparisons across groups were performed with the Kruskal-Wallis test. Resulting significance was followed up with post-hoc tests between groups with Wilcoxon signed-rank test for paired data and Mann Whitney U test for un-paired data. When comparing groups containing normal and non-normal data, non-parametric tests were applied.

When comparing the relationship between gait features to speed, the Pearson correlation coefficient was calculated for reach length, step length, step width, reach duration, and duty factor. The Spearman correlation coefficient was calculated for grasp duration given its nonlinear relationship with speed.

Circular data (i.e. phase offset values for coordination analysis, and angles for trajectory analysis) were analyzed using appropriate circular counterparts to standard descriptive statistics measures and statistical tests with the *pycircstat2* toolbox in Python to account for circular affect. Rayleigh’s r-test was used to determine if circular distributions were non-uniform (p < 0.05) and to identify the mean resultant vector, which indicates how disperse the data are with respect to the mean. Watson-Williams test was used to determine if there were significant differences across multiple groups of phase offset values for different paw pairings.

In all figures, the following convention was used: *p < 0.05, **p < 0.001, and n.s. for non-significant. When appropriate, multiple comparisons were tested for using the Bonferroni correction to adjust p-values for significance.

## Acknowledgements

Thank you to Dr. Laura Andreoli and Antonia Constantinescu from University College London for discussions on behavioral design and analysis. This work was supported by a Brain Research United Kingdom fellowship (to C.J.B.), and a Medical Research Foundation Fellowship MRF-087-0003-F-KOCH-C0917 (to S.C.K.).

## Author Contributions

Conceptualization, C.J.B; Data curation, C.J.B; Formal analysis, C.J.B; Investigation, C.J.B.; Methodology, C.J.B. Project administration, C.J.B., L.E.B., R.M.B., S.C.K.; Software, C.J.B.; Resources, C.J.B., L.E.B., R.M.B., S.C.K.; Validation, C.J.B. Visualization, C.J.B. Writing – original draft, C.J.B.; Writing – reviewing and editing, C.J.B., L.E.B., R.M.B, S.C.K.

## Declarations

The authors declare no competing interests.

## Data availability

Data will be made available following reasonable request to S.C.K. Pertinent files for wall construction are available at https://github.com/cjblack/TheWall.

## Supplemental

**Figure S1.**
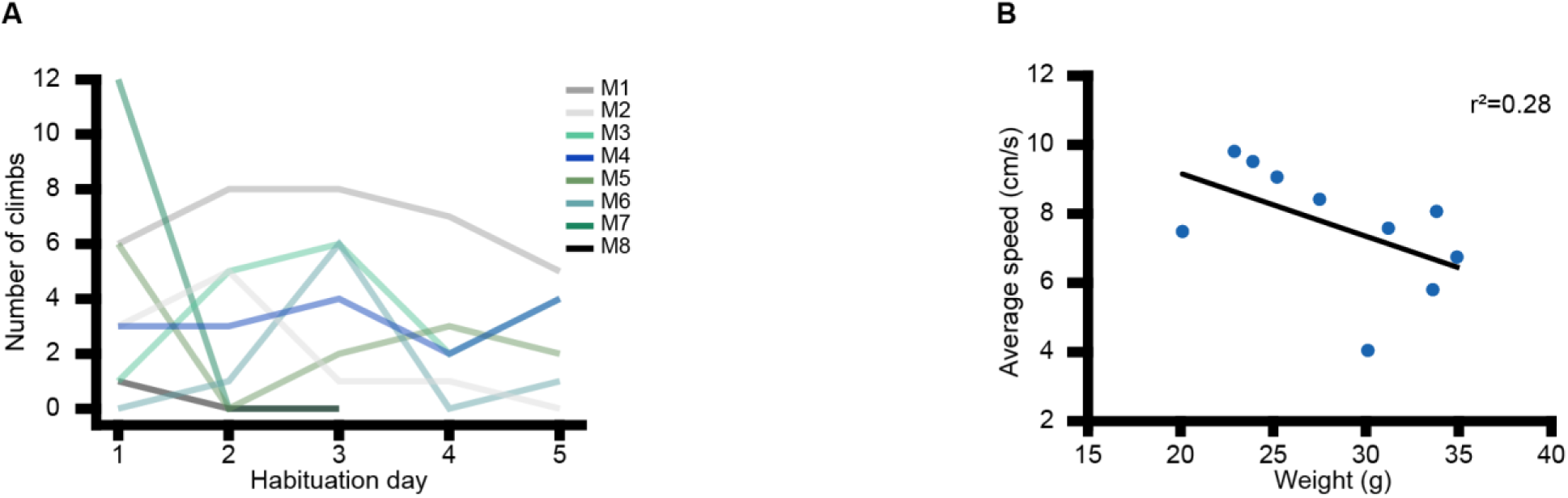
Habituation and weight. (A) Spontaneous climbs across habituation days for each mouse; only two mice did not spontaneously climb during habituation (not shown in figure). (B) Climbing speed as a function of weight, linear regression shows a negative trend indicating slower average speed was related to body weight.

**Figure S2.**
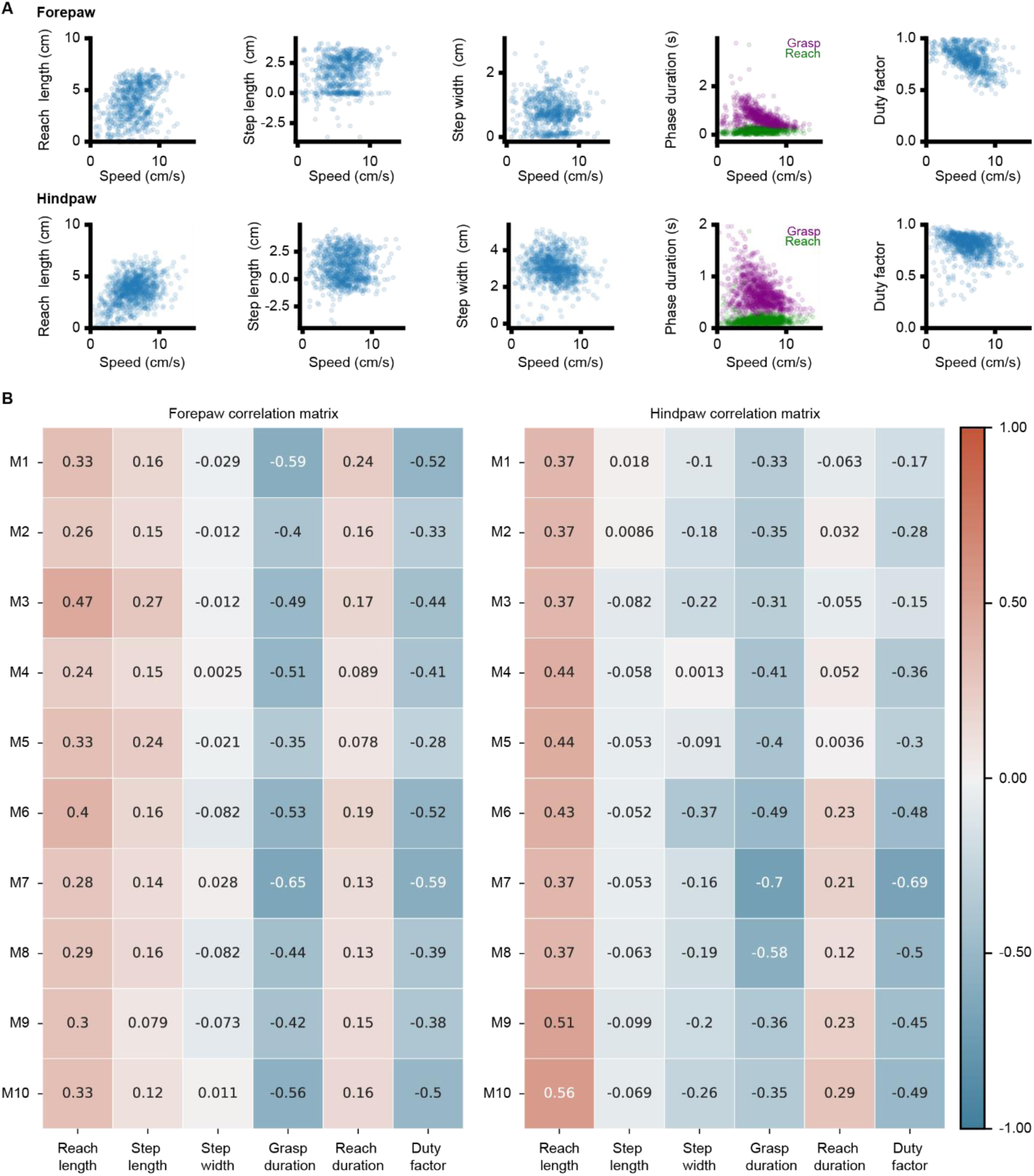
Correlations between speed and kinematic variables. (A) Example forepaw (top row) and hindpaw (bottom row) scatter plots showing relationship between speed and reach length, step length, step width, phase duration, and duty factor from one mouse. (B) Correlations for speed and kinematic variables for all mice (M1-M10) for forepaw (left) and hindpaw (right) data.

**Table S1.**
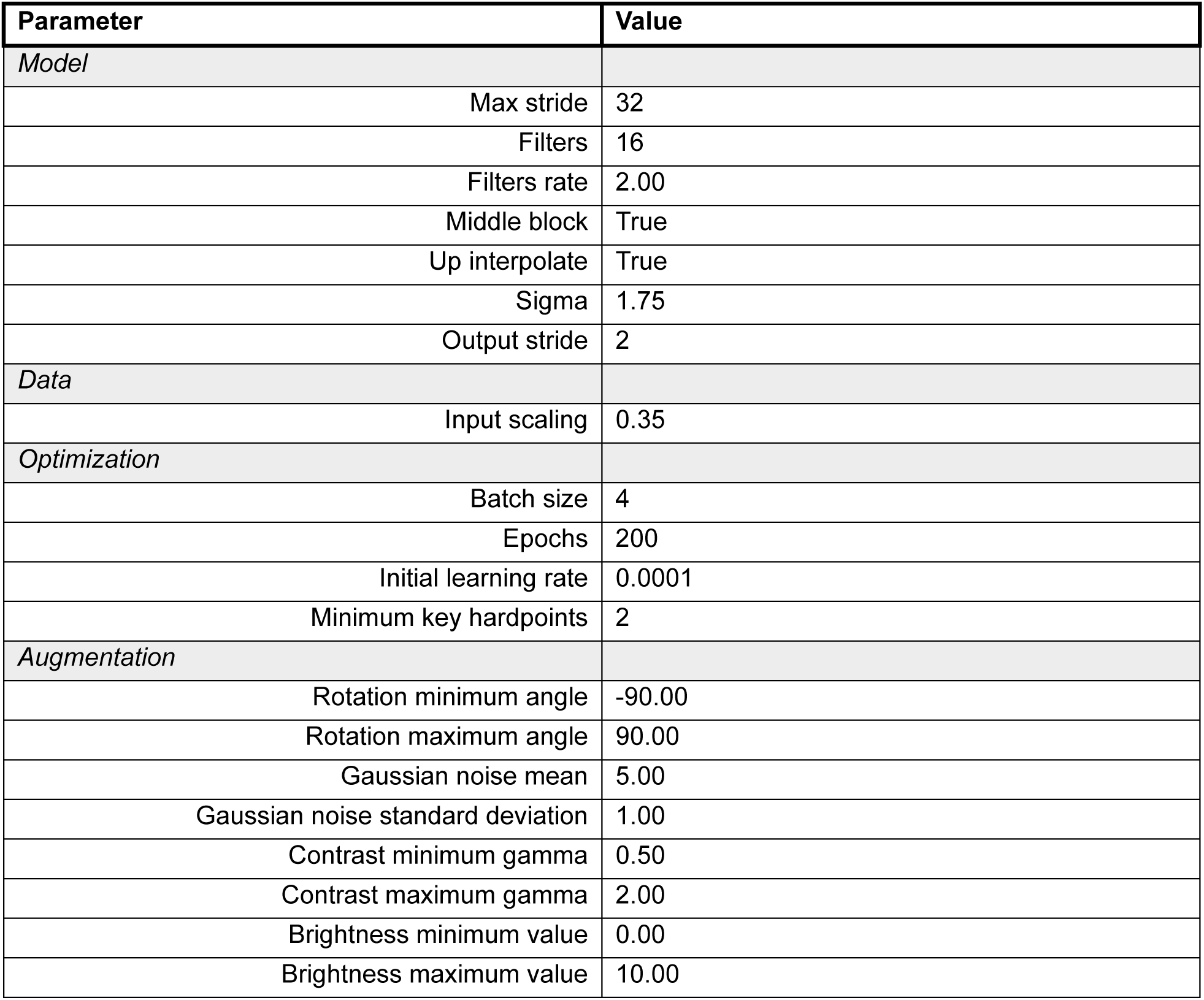
Network hyperparameters for pose estimation.

## References

1. Flash, T., & Hochner, B. (2005). Motor primitives in vertebrates and invertebrates. Current opinion in neurobiology, 15(6), 660–666. 10.1016/j.conb.2005.10.011.

2. Giszter, S. F. (2015). Motor primitives—new data and future questions. Current opinion in neurobiology, 33, 156–165.10.1016/j.conb.2015.04.004.

3. Grafton, S. T. (2010). The cognitive neuroscience of prehension: recent developments. Experimental brain research, 204(4), 475–491. 10.1007/s00221-010-2315-2.

4. Mathis, M.W., Mathis, A., and Uchida, N. (2017). Somatosensory Cortex Plays an Essential Role in Forelimb Motor Adaptation in Mice. Neuron 93, 1493–1503.e6. 10.1016/j.neuron.2017.02.049.

5. Mosberger, A.C., Sibener, L.J., Chen, T.X., Rodrigues, H.F.M., Hormigo, R., Ingram, J.N., Athalye, V.R., Tabachnik, T., Wolpert, D.M., Murray, J.M., et al. (2024). Exploration biases forelimb reaching strategies. Cell Rep 43. 10.1016/j.celrep.2024.113958.

6. Donegan, D., Kanzler, C.M., Büscher, J., Viskaitis, P., Bracey, E.F., Lambercy, O., and Burdakov, D. (2022). Hypothalamic Control of Forelimb Motor Adaptation. Journal of Neuroscience 42, 6243–6257. 10.1523/JNEUROSCI.0705-22.2022.

7. Galiñanes, G.L., Bonardi, C., and Huber, D. (2018). Directional Reaching for Water as a Cortex-Dependent Behavioral Framework for Mice. Cell Rep 22, 2767–2783. 10.1016/j.celrep.2018.02.042.

8. Vergara-Aragon, P., Gonzalez, C.L.R., and Whishaw, I.Q. (2003). A novel skilled-reaching impairment in paw supination on the “good” side of the hemi-Parkinson rat improved with rehabilitation. Journal of Neuroscience 23, 579–586. 10.1523/jneurosci.23-02-00579.2003.

9. 9. Khademullah, C.S., and De Koninck, Y. (2022). A novel assessment of fine-motor function reveals early hindlimb and detectable forelimb deficits in an experimental model of ALS. Sci Rep 12. 10.1038/s41598-022-20333-1.

10. Woodard, C.L., Sepers, M.D., and Raymond, L.A. (2021). Impaired refinement of kinematic variability in huntington disease mice on an automated home cage forelimb motor task. Journal of Neuroscience 41, 8589–8602. 10.1523/JNEUROSCI.0165-21.2021.

11. Karantanis, N.E., Rychlik, L., Herrel, A., and Youlatos, D. (2018). Vertical Locomotion in Micromys minutus (Rodentia: Muridae): Insights into the Evolution of Eutherian Climbing. J Mamm Evol 25, 277–289. https://doi.org/10.1007/s10914-016-9374-5.

12. Seifert, L., Orth, D., Mantel, B., Boulanger, J., Hérault, R., and Dicks, M. (2018). Affordance realization in climbing: Learning and transfer. Front Psychol 9. 10.3389/fpsyg.2018.00820.

13. Ishiwaka, R., and Möri, T. (1999). Early development of climbing skills in harvest mice. Anim Behav 58, 203–209. 10.1006/anbe.1999.1146.

14. Bourdin, C., Teasdale, N., Nougier, V., Bard, C., and Fleury, M. (1999). Postural constraints modify the organization of grasping movements. Hum Mov Sci 18, 87–102. 10.1016/S0167-9457(98)00036-0.

15. Bhowmick, S., D‘Mello, V., Ponery, N., and Abdul-Muneer, P.M. (2018). Neurodegeneration and sensorimotor deficits in the mouse model of traumatic brain injury. Brain Sci 8. 10.3390/brainsci8010011.

16. Santos, E.J., Giddings, A.N., Kandil, F.A., and Negus, S.S. (2023). Climbing behavior by mice as an endpoint for preclinical assessment of drug effects in the absence and presence of pain. Frontiers in pain research (Lausanne, Switzerland) 4, 1150236. 10.3389/fpain.2023.1150236.

17. Kaltenbach, L.S., Romero, E., Becklin, R.R., Chettier, R., Bell, R., Phansalkar, A., Strand, A., Torcassi, C., Savage, J., Hurlburt, A., et al. (2007). Huntingtin interacting proteins are genetic modifiers of neurodegeneration. PLoS Genet 3, 689–708. 10.1371/journal.pgen.0030082.

18. Pereira, T.D., Tabris, N., Matsliah, A., Turner, D.M., Li, J., Ravindranath, S., Papadoyannis, E.S., Normand, E., Deutsch, D.S., Wang, Z.Y., et al. (2022). SLEAP: A deep learning system for multi-animal pose tracking. Nat Methods 19, 486–495. 10.1038/s41592-022-01426-1.

19. Lopes, G., Bonacchi, N., Frazão, J., Neto, J.P., Atallah, B. V., Soares, S., Moreira, L., Matias, S., Itskov, P.M., Correia, P.A., et al. (2015). Bonsai: An event-based framework for processing and controlling data streams. Front Neuroinform 9. 10.3389/fninf.2015.00007.

20. Bains, R.S., Forrest, H., Sillito, R.R., Armstrong, J.D., Stewart, M., Nolan, P.M., and Wells, S.E. (2023). Longitudinal home-cage automated assessment of climbing behavior shows sexual dimorphism and aging-related decrease in C57BL/6J healthy mice and allows early detection of motor impairment in the N171-82Q mouse model of Huntington’s disease. Front Behav Neurosci 17. 10.3389/fnbeh.2023.1148172.

21. Chen, C.-C., Gilmore, A., and Zuo, Y. (2014). Study motor skill learning by single-pellet reaching tasks in mice. J Vis Exp. 10.3791/51238.

22. Burnside, E.R., De Winter, F., Didangelos, A., James, N.D., Andreica, E.C., Layard-Horsfall, H., Muir, E.M., Verhaagen, J., and Bradbury, E.J. (2018). Immune-evasive gene switch enables regulated delivery of chondroitinase after spinal cord injury. Brain 141, 2362–2381. 10.1093/brain/awy158.

23. Hruska, R.E., Kennedy, S., and Silbergeld, E.K. (1979). Quantitative aspects of normal locomotion in rats. Life Sci 25, 171–179. 10.1016/0024-3205(79)90389-8.

24. Heglund, N.C., Taylor, C.R., and Mcmahon, T.A. (1974). Scaling stride frequency and gait to animal size: Mice to horses. Science (1979) 186, 1112–1113. 10.1126/science.186.4169.1112.

25. Grillner, S., Halbertsma, J., Nilsson, J., and Thorstensson, A. (1979). The adaptation to speed in human locomotion. Brain Res 165, 177–182. 10.1016/0006-8993(79)90059-3.

26. Frigon, A. (2017). The neural control of interlimb coordination during mammalian locomotion. J Neurophysiol 117, 2224–2241. 10.1152/jn.00978.2016.

27. Milner, T.E., and Ijaz, M.M. (1990). The effect of accuracy constraints on three-dimensional movement kinematics. Neuroscience 35, 365–374. 10.1016/0306-4522(90)90090-Q.

28. Becker, M.I., Calame, D.J., Wrobel, J., and Person, A.L. (2020). Online control of reach accuracy in mice. J Neurophysiol 124, 1637–1655. 10.1152/jn.00324.2020.

29. Marteniuk, R.G., MacKenzie, C.L., Jeannerod, M., Athenes, S., and Dugas, C. (1987). Constraints on human arm movement trajectories. Can J Psychol 41, 365–378. 10.1037/h0084157.

30. Todorov, E., and Jordan, M.I. (1998). Smoothness maximization along a predefined path accurately predicts the speed profiles of complex arm movements. J Neurophysiol 80, 696–714. 10.1152/jn.1998.80.2.696.

31. Celikel, T., and Sakmann, B. (2007). Sensory integration across space and in time for decision making in the somatosensory system of rodents. Proc Natl Acad Sci U S A 104, 1395–1400. 10.1073/pnas.0610267104.

32. Voigts, J., Sakmann, B., and Celike, T. (2008). Unsupervised whisker tracking in unrestrained behaving animals. J Neurophysiol 100, 504–515. 10.1152/jn.00012.2008.

33. 33. Zong, W., Obenhaus, H.A., Skytøen, E.R., Eneqvist, H., de Jong, N.L., Vale, R., Jorge, M.R., Moser, M.B., and Moser, E.I. (2022). Large-scale two-photon calcium imaging in freely moving mice. Cell 185, 1240–1256.e30. 10.1016/j.cell.2022.02.017.

34. 34. Peris-Sampedro, F., Stoltenborg, I., Le May, M. V., Zigman, J.M., Adan, R.A.H., and Dickson, S.L. (2021). Genetic deletion of the ghrelin receptor (GHSR) impairs growth and blunts endocrine response to fasting in Ghsr-IRES-Cre mice. Mol Metab 51. 10.1016/j.molmet.2021.101223.

35. Sober, S.J., and Sabes, P.N. (2005). Flexible strategies for sensory integration during motor planning. Nat Neurosci 8, 490–497. 10.1038/nn1427.

36. Klein, L.K., Maiello, G., Stubbs, K., Proklova, D., Chen, J., Paulun, V.C., Culham, J.C., and Fleming, R.W. (2023). Distinct neural components of visually guided grasping during planning and execution. Journal of Neuroscience 43, 8504–8514. 10.1523/JNEUROSCI.0335-23.2023.

37. Liu, D., and Todorov, E. (2007). Evidence for the flexible sensorimotor strategies predicted by optimal feedback control. Journal of Neuroscience 27, 9354–9368. 10.1523/JNEUROSCI.1110-06.2007.

38. Betti, S., Castiello, U., & Begliomini, C. (2021). Reach-to-grasp: a multisensory experience. Frontiers in Psychology, 12, 614471 10.3389/fpsyg.2021.614471.

39. Parmiani, P., Lucchetti, C., and Franchi, G. (2018). Whisker and nose tactile sense guide rat behavior in a skilled reaching task. Front Behav Neurosci 12. 10.3389/fnbeh.2018.00024.

40. Sartori, L., Camperio-Ciani, A., Bulgheroni, M., and Castiello, U. (2014). How posture affects macaques’ reach-to-grasp movements. Exp Brain Res 232, 919– 925. 10.1007/s00221-013-3804-x.

41. Kaminski, T.R., and Simpkins, S. (2001). The effects of stance configuration and target distance on reaching: I. Movement preparation. Exp Brain Res 137, 439–446. 10.1007/s002210000604.

42. Warren, R.A., Zhang, Q., Hoffman, J.R., Li, E.Y., Hong, Y.K., Bruno, R.M., and Sawtell, N.B. (2021). A rapid whisker-based decision underlying skilled locomotion in mice. Elife 10. 10.7554/eLife.63596.

43. Green, D.J., Richmond, B.G., and Miran, S.L. (2012). Mouse Shoulder Morphology Responds to Locomotor Activity and the Kinematic Differences of Climbing and Running. J Exp Zool B Mol Dev Evol 318, 621–638. 10.1002/jez.b.22466.

44. Nirody, J.A., Duran, L.A., Johnston, D., and Cohen, D.J. (2021). Tardigrades exhibit robust interlimb coordination across walking speeds and terrains. Proc Natl Acad Sci U S A 118. 10.1073/pnas.2107289118.

